# Neurons expressing pathological Tau protein trigger dramatic changes in microglial morphology and dynamics

**DOI:** 10.1101/731901

**Authors:** Rahma Hassan-Abdi, Alexandre Brenet, Mohamed Bennis, Constantin Yanicostas, Nadia Soussi-Yanicostas

## Abstract

Microglial cells, the resident macrophages of the brain, are important players in the pathological process of numerous neurodegenerative disorders, including tauopathies, a heterogeneous class of diseases characterized by intraneuronal Tau aggregates. However, microglia response in Tau pathologies remains poorly understood. Here we exploit a genetic zebrafish model of tauopathy, combined with live microglia imaging, to investigate the behaviour of microglia *in vivo* in the disease context. Results show that while microglia were almost immobile and displayed long and highly dynamic branches in a wild-type context, in presence of diseased neurons cells became highly mobile and displayed morphological changes, with highly mobile cell bodies together with fewer and shorter processes. We also imaged, for the first time to our knowledge, the phagocytosis of apoptotic tauopathic neurons by microglia *in vivo* and observed that microglia engulfed about as twice materials as in controls. Finally, genetic ablation of microglia in zebrafish tauopathy model significantly increased Tau hyperphosphorylation, suggesting that microglia provide neuroprotection to diseased neurons. Our findings demonstrate for the first time the dynamics of microglia in contact with tauopathic neurons *in vivo* and open perspectives for the real-time study of microglia in many neuronal diseases.

## 1 Introduction

Microglia, the resident brain macrophages, are highly plastic and multifunctional cells that continuously monitor the health of neuronal networks (Kierdorf and Prinz, 2017). In a physiological context, microglia display long cytoplasmic processes that constantly extend and retract to contact neighbour neurons and check their physiology (Nimmerjahn et al., 2005; Peri and Nüsslein-Volhard, 2008). Microglia also respond promptly to brain injury or infection, with both immuno-protective and cytotoxic responses, including the secretion of a large set of cytokines (Butovsky and Weiner, 2018; Hanisch, 2002; Hu et al., 2015; Wake et al., 2013) and increased phagocytic capacities to eliminate pathogen debris and dead cells (Leong and Ling, 1992; Ling and Wong, 1993; Brockhaus et al., 1996; Nakajima and Kohsaka, 2001; Hanisch and Kettenmann, 2007; Thameem Dheen et al., 2007). However, in some disease contexts, such as tauopathies, microglia also appear to have harmful activities (Bhaskar et al., 2010; Eyo and Dailey, 2013; Maphis et al., 2015; Laurent et al., 2018).

Tauopathies are a family of neurodegenerative disorders characterized by intra-neuronal fibrillary aggregates containing abnormally hyperphosphorylated isoforms of the microtubule-associated protein Tau (Alavi Naini and Soussi-Yanicostas, 2015; Spillantini and Goedert, 2013; Wang and Mandelkow, 2016). While the causal role of Tau in the disease is supported by several inherited tauopathies triggered by dominant missense mutations in the protein, such as Tau^P301L^, causing fronto-temporal dementia with parkinsonism on chromosome 17 (FTDP-17) (Hutton et al., 1998), the aetiology of these disorders and the contribution of microglia to their physiopathology remain poorly understood (Hansen et al., 2018; Laurent et al., 2018; Perea et al., 2018).

Because of their plasticity and well-established neuroprotective activities, microglial cells are very promising therapeutic targets for the treatment of neuron disorders, including neurodegenerative diseases.

In an attempt to describe the behavior of microglial cells in a tauopathy disease context *in vivo*, we used the transgenic zebrafish Tg(HuC-hTau^P301L^:DsRed) tauopathy model (Paquet al., 2009) and live microglia imaging (Peri and Nüsslein-Volhard, 2008). We observed that in the presence of hTau^P301L^-expressing neurons, microglia display dramatic changes in morphology and dynamics, with cells showing fewer and shorter branches and amoeboid-like cell bodies alongside a markedly increased mobility and phagocytic activity. We also imaged the phagocytosis of dying neurons by microglia and showed that these cells could phagocyte nearly twice as much as in homeostatic brains. However, we also observed that these microglial cells failed to phagocyte all dead neurons, highlighting the limits of their phagocyting abilities.

## 2 Results

### 2.1 Microglia display dramatic changes in shape and dynamics in the presence of hTau^P301L^-expressing neurons

To investigate the behavior of microglial cells in a tauopathy disease context *in vivo*, we used the transgenic Tg(HuC-hTau^P301L^:DsRed) zebrafish model of Tau-induced neurodegeneration, combined with the transgenic Tg(ApoE-eGFP) microglia marker line. As previously shown, in the optic tectum of Tg(ApoE-eGFP) embryos, microglia displayed a ramified morphology, with a small cell body and several elongated branches (Figures 1A,C). By contrast, in Tg(ApoE-eGFP; HuC-hTau^P301L^:DsRed) embryo microglia displayed a rounder morphology, with a larger cell body and fewer, shorter branches (Figures 1B,B’,D). Quantifications of morphological parameters confirmed these dramatic changes in microglia morphology seen in the presence of diseased neurons, with a smaller surface area (Figure 1E) and volume (Figure 1F); and a greater sphericity (Figure 1G). However, alongside these rounded microglia, a few branched cells were also observed in the disease context (Figures 1B, B’).

**Figure 1.**
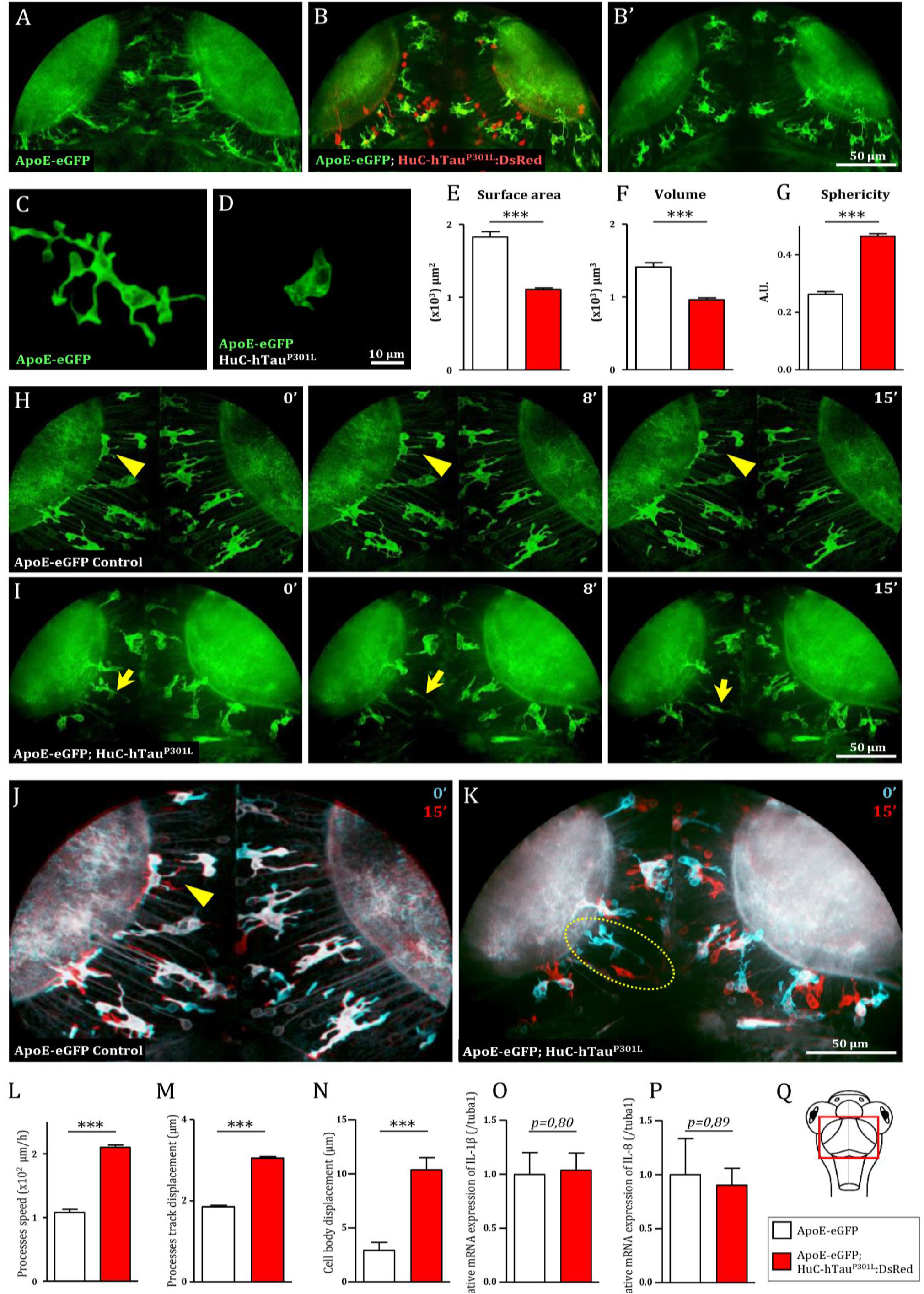
Microglia displays dramatic changes in morphology and dynamics in the presence of hTau^P301L^ –expressing neurons. (**A, B, B’**) Dorsal views of the optic tectum of 7 dpf Tg(ApoE-eGFP) (**A**) and Tg(ApoE-eGFP; HuC-hTau^P301L^:DsRed) transgenic embryos (**B, B’**), showed the characteristic ramified morphology of microglia in wild-type (**A**), while in the presence of hTau^P301L^-expressing neurons, microglial cells displayed shorter processes and larger cell bodies. (**C, D**) Detailed morphology of microglial cells in Tg(ApoE-eGFP) (**C**) and Tg(ApoE-eGFP; HuC-hTau^P301L^:DsRed) embryos (**D**). (**E-G**) Measurements of microglia morphological parameters; surface area (**E**, *p* < 0.0001), volume (**F**, *p* < 0.0001) and sphericity (**G**, *p* < 0.0001), in Tg(ApoE-eGFP) (n=10) and Tg(ApoE-eGFP; HuC-hTau^P301L^:DsRed) (n=24) embryos, confirmed the cell shape changes observed in the presence of hTau^P301L^-expressing neurons. (**H, I**) Time-lapse sequences of microglia dynamics in Tg(ApoE-eGFP) (**H**, Video 1) and Tg(ApoE-eGFP; HuC-hTau^P301L^:DsRed) embryos (**I**, Video 2). (**J, K**) Merged images of two time points separated by 15 minutes from video 1 (**J**) and video 2 (**K**). The merged images at t=0 minutes (cyan) and t=15 minutes (red) highlighted the dramatic increased mobility of microglial cell bodies in the presence of hTau^P301L^-expressing neurons. (**L-N**) Measurements of microglia dynamics; process speed (**L**, *p* = 0.0004), process track displacement (**M**, *p* = 0.0002) and cell body displacement (**N**, *p* = 0.0054), in Tg(ApoE-eGFP) (n=3) and Tg(ApoE-eGFP; HuC-hTau^P301L^:DsRed) (n=4) embryos, confirmed the increased mobility of both microglia processes and cell bodies observed in the presence of hTau^P301L^-expressing neurons. (**O, P**) Measurements of pro-inflammatory cytokine expression in the brain of 5 dpf Tg(ApoE-eGFP) (n=6) and Tg(ApoE-eGFP; HuC-hTau^P301L^:DsRed) (n=11) embryos. Comparison of the relative expression of IL-1β (**O**, *p* = 0.80) and IL-8 (**P**, *p* = 0.89) in both groups shows no significant differences. (**Q**) Schematic dorsal view of a 7 dpf zebrafish embryo. The red square shows the region of interest that comprises the optic tectum. ***: *p* < 0.001; **: *p* < 0.01; *= *p* < 0.05. Scale bar (A, B, B’, H, I, J, K) = 50 µm, (C, D) = 10 µm. A.U.: arbitrary units.

Given that microglial cells are highly dynamic, we used *in vivo* real-time confocal imaging combined with Imaris software (Bitplane Inc.) image analysis to determine whether the presence of hTau^P301L^-expressing neurons modified microglia dynamics. In Tg(ApoE-eGFP) embryos, microglia displayed dynamic processes that were constantly extending and retracting, while their cell bodies remained almost immobile (Figures 1H,J, Video 1, Supplementary figures 1A,C, Video 5). By contrast, in Tg(ApoE-eGFP; HuC-hTau^P301L^:DsRed) embryos, microglia were highly mobile with their cell bodies traveling over longer distances (Figures 1I,K, Video 2, Supplementary figures 1B,D, Video 6). Quantifications of microglia dynamics confirmed that in the presence of hTau^P301L^-expressing neurons, microglia displayed increased mean process speed (Figure 1L) and mean process track displacement (Figure 1M), and a much larger displacement of the cell bodies over a similar time frame (Figure 1N).

To further characterize the phenotype of microglial cells exposed to hTau^P301L^-expressing neurons, we analysed the expression levels of the pro-inflammatory cytokines, IL-1β, IL-8 and TNF-α in the brain tissue of transgenic Tg(HuC-hTau^P301L^:DsRed) and wild-type embryos. Unexpectedly, none of these cytokines were overexpressed in the pathologic context, the two tested groups displaying no significant differences in expression levels of IL-1β, IL-8 (Figures 1O,P) and TNF-α (data not shown).

### 2.2 Genetic depletion of microglia worsens pathology in Tg(HuC-hTau^P301L^:DsRed) embryos

As a first attempt to investigate the function of microglial cells in Tau pathology, we generated Tg(HuC-hTau^P301L^:DsRed) embryos completely devoid of microglia following injection of an antisense morpholino oligonucleotide targeting pU.1 (MO-pU1) transcripts encoding a transcription factor essential for proper differentiation of macrophage/microglia (Rhodes et al., 2005), and then studied the consequences of such microglial cell ablation on Tau phosphorylation, neuron apoptosis and expression of pro-inflammatory cytokines. Injection of the MO-pU1 (Figure 2A) leads to a complete absence of microglial cells in the brain of the embryos as shown by either Neutral Red staining (Figure 2B), or immunocytochemistry using L-plastin antibody (Figure 2C).

**Figure 2.**
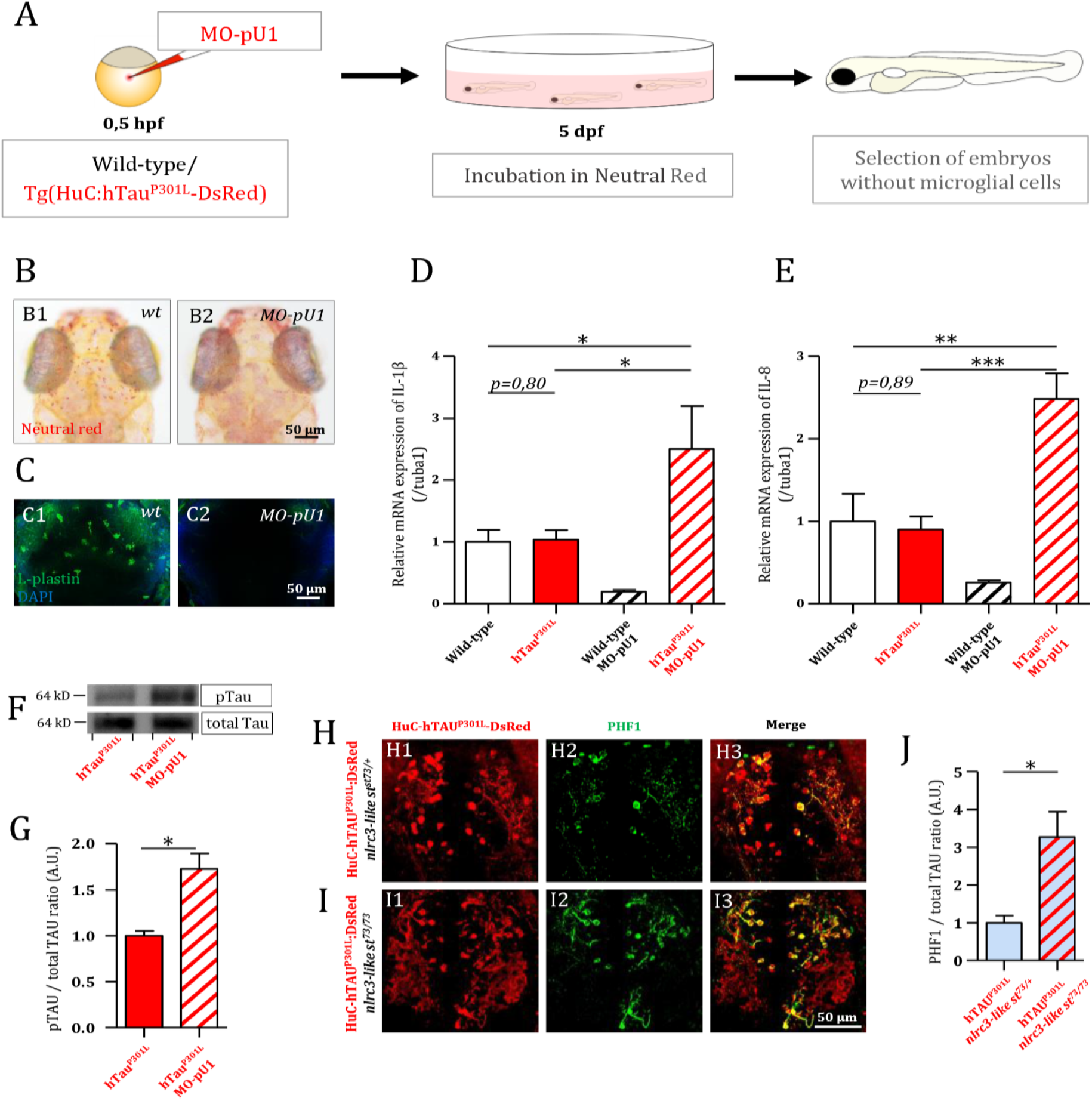
Genetic depletion of microglia worsens the pathology in Tg(HuC-hTau^P301L^:DsRed) embryos. (**A**) Outline of microglia depletion experiments. Embryos were injected at the single cell stage with a solution of antisense morpholino oligonucleotide targeting pU.1 transcripts. At 5 dpf, injected embryos were incubated in Neutral Red solution to sort microglia-depleted embryos. (**B**) Dorsal views of the optic tectum of 5 dpf wild-type microglia-depleted (**B2**) and untreated live embryos (**B1**), following incubation in Neutral Red solution. (**C**) Dorsal views of the optic tectum of 5 dpf wild-type microglia-depleted (**C2**) and untreated fixed embryos (**C1**), labelled with L-plastin antibody. (**D**, **E**) Measurements of pro-inflammatory cytokines in the brain of 5 dpf wild-type embryos with (n=6), or without (n=3) microglia; and Tg(HuC-hTau^P301L^:DsRed) embryos with (n=11), or without microglia (n=7). Both relative expressions of IL-1β (**D**, *p* = 0.035) and IL-8 (**E**, *p* < 0.0001) display a significant increase in the brains of Tg(HuC-hTau^P301L^:DsRed) embryos without microglia cells, compared to their siblings with microglial cells. (**F**, **G**) Representative Western blots membranes of total protein extracts from 6 dpf Tg(HuC-hTau^P301L^:DsRed) embryos with (left) or without (right) microglia, hybridized with antibodies against human total Tau (total Tau) or human phosphorylated Tau at Ser396 residue (pTau) (**F**); and quantification of corresponding pTau/total Tau ratio (respectively, n=4 and n=4) (**G**, *p* = 0.01). The ratio of hyperphosphorylated hTau to total Tau protein is significantly increased in microglia-depleted Tg(HuC-hTau^P301L^:DsRed) embryos. (**H**-**J**) Dorsal views of the telencephalon of 6 dpf Tg(HuC-hTau^P301L^:DsRed; *nlrc3-like^st73/+^*) embryos (**H**) and Tg(HuC-hTau^P301L^:DsRed; *nlrc3-like^st73/73^*) embryos (**I**), labelled with an antibody directed against human phosphorylated Tau at Ser396 and Ser404 residues (PHF1); and quantification of corresponding PHF1/hTau^P301L^-DsRed signal ratio (respectively, n=4 and n=6) (**J**, *p* = 0.0485). The quantification of the signal ratio of hyperphosphorylated hTau protein on brain sections from Tg(HuC-hTau^P301L^:DsRed; *nlrc3-like^st73/73^*) mutant embryos devoid of microglia confirmed the significant increase of this ratio displayed in protein extracts from Tg(HuC-hTau^P301L^:DsRed) embryos microglia-depleted with morpholino. ***, *p* < 0.001; **, *p* < 0.01; *, *p* < 0.05. Scale bar (B, C, H, I) = 50 µm.

Using 5 dpf wild-type and transgenic Tg(HuC-hTau^P301L^:DsRed) embryos and microglia ablation following MO-pU1 injection, we first studied the consequences of the absence of microglia on the expression of pro-inflammatory cytokines IL-1β and IL-8. Results showed that while expression of Tau^P301L^ did not stimulate overexpression of IL-1β (Figure 2D) and IL-8 (Figure 2E) in embryos with microglia embryos, microglia depletion in Tg(HuC-hTau^P301L^:DsRed) embryos provoked a markedly increased expression of both these pro-inflammatory cytokines.

As a first attempt to determine the effect of the absence of microglia on Tau hyperphosphorylation *in vivo*, we quantified and compared hTau phosphorylation levels at Ser396 site in Tg(HuC-hTau^P301L^:DsRed) embryos with and without microglia (Figure 2F). Interestingly, in Tg(HuC-hTau^P301L^:DsRed) embryos without microglia, we observed an increased accumulation of hyperphosphorylated Tau when compared to that seen in their siblings with microglia (Figure 2F).

Quantification of phospho-Tau to total Tau accumulation ratio (pTau/Tau) confirmed that hTau hyperphosphorylation levels were significantly increased in microglia-depleted Tg(HuC-hTau^P301L^:DsRed) embryos (Figure 2G). To further investigate the consequences of the absence of microglia on Tau hyperphosphorylation, Tg(HuC-hTau^P301L^:DsRed) mutant embryos, which are fully devoid of microglia as the result of homozygous nlrc3-like^st73^ mutation (Shiau et al., 2013), and analyzed hTau^P301L^ hyperphosphorylation using the antibody PHF1, targeting pathological phosphorylation sites Ser396 and Ser404 of the hTau protein (Figures 2H,I). In good agreement with Western blot analysis, a significant increase in PHF1 labelling intensity was observed in the telencephalon of 6 dpf Tg(HuC-hTau^P301L^:DsRed; *nlrc3-like^st73/73^*) mutant embryos (Figure 2I) when compared to that observed in the brain of their Tg(HuC-hTau^P301L^:DsRed; *nlrc3-like^st73/+^*) siblings with microglia (Figure 2H). Quantification of the signal ratio of hyperphosphorylated hTau protein on brain sections from Tg(HuC-hTau^P301L^:DsRed; *nlrc3-like^st73/73^*) embryos confirmed the significant increase of this ratio displayed in protein extracts from Tg(HuC-hTau^P301L^:DsRed) embryos microglia-depleted with morpholino (Figure 2J).

### 2.3 Microglia phagocytic activity is enhanced in the presence of hTau^P301L^-expressing neurons

As phagocytosis is a main feature of microglial cells, we first monitored the phagocytic activity of microglia in Tg(ApoE-eGFP; HuC-hTau^P301L^:DsRed) embryos. We observed the phagocytosis of hTau^P301L^-expressing neurons by microglia, using confocal real-time imaging (Figure 3B, Video 3). A microglial cell in the optic tectum (Figures 3B,C,0 min) sends one of its processes to the pathological neuron (Figures 3B,C,5 min) to draw it towards its cell body (Figures 3B,C,9 min) and execute the digestion of the neuron and its debris until completion of the process (Figures 3B,C,18 min). We also observed the detail of a microglial cell engulfing three neurons simultaneously (Supplementary Figure 2, Video 7). We next assessed the phagocytic activity of microglia by quantifying the total engulfed volume, which was significantly increased in Tg(ApoE-eGFP; HuC-hTau^P301L^:DsRed) embryos (Figure 3D). Given the critical role of microglia in removing apoptotic cells and other noxious elements, we next visualized neuronal death in Tg(ApoE-eGFP; HuC-hTau^P301L^:DsRed) embryos using the apoptotic marker Acridine Orange. Data showed that microglia specifically engulfed apoptotic neurons (Figures 3E-F, Video 4) but not non-apoptotic hTau^P301L^-expressing cells, supporting the notion that microglia specifically responds to signals sent by degenerating neurons that are already apoptotic but not hTau^P301L^-expressing neurons *per se*. However, quantification of the number of non-engulfed apoptotic neurons in Tg(ApoE-eGFP; HuC-hTau^P301L^:DsRed) and control Tg(ApoE-eGFP; HuC-RFP) embryos showed that microglia failed to phagocyte all apoptotic hTau^P301L^-expressing neurons (Figure 3K).

**Figure 3.**
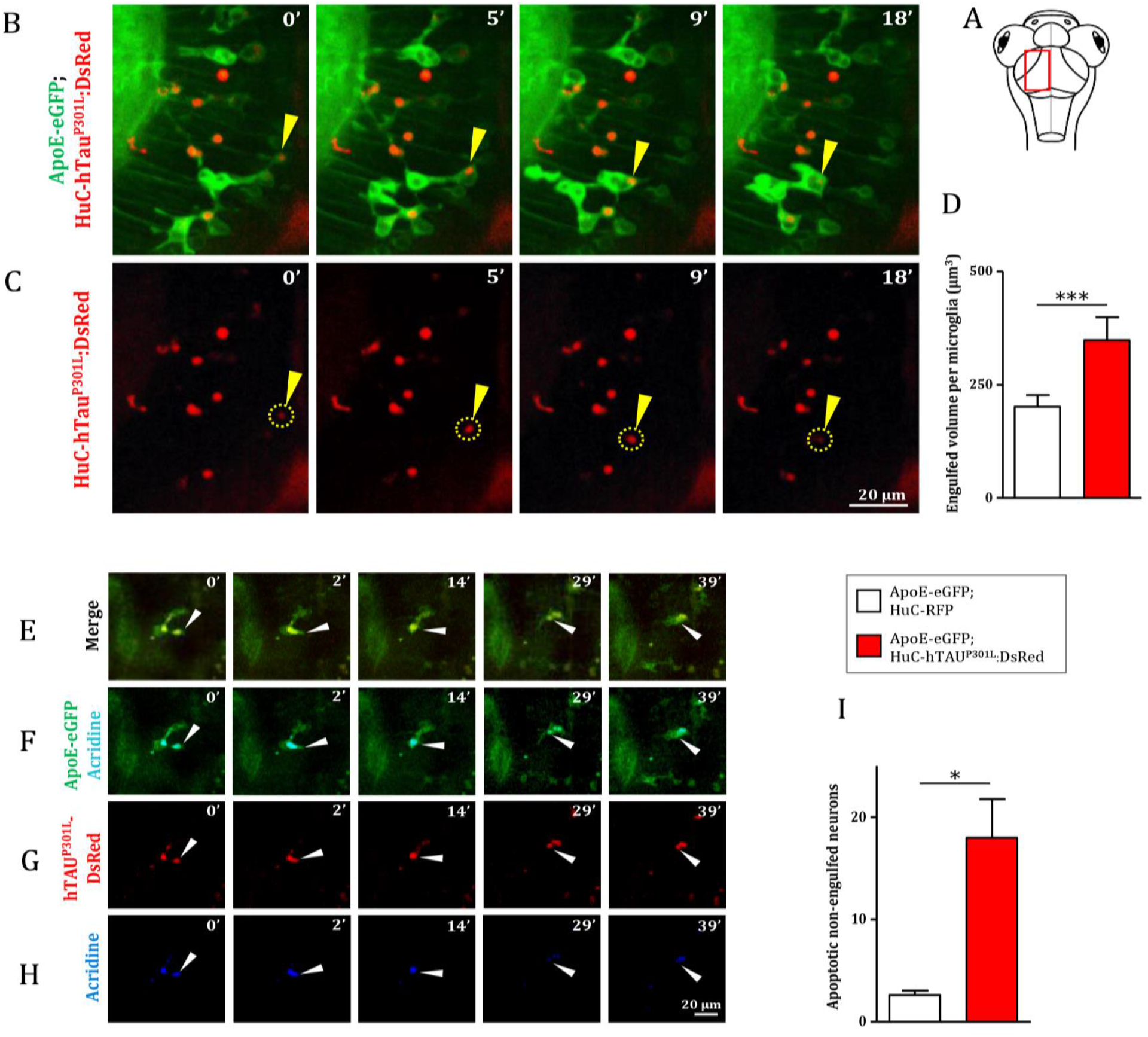
Microglia phagocytic activity is increased in presence of hTau^P301L^–expressing, but appears non-sufficient in eliminating all apoptotic neurons. (**A**) Schematic illustration of 7 dpf embryo in dorsal view. The red square shows the region of the optic tectum where the time-lapse (**B, C**) was recorded. (**B, C**) Time-lapse imaging of a microglial cell phagocyting a diseased neuron (yellow arrowhead) in a 7 dpf Tg(ApoE-eGFP; HuC-hTau^P301L^:DsRed) embryo; (**B**, Video 3) merge of GFP and DsRed; (**C**) DsRed only. (**D**, *p* = 0.0262) Quantification of the engulfed neuronal volume in Tg(ApoE-eGFP; HuC-RFP) (n=7) and Tg(ApoE-eGFP; HuC-hTau^P301L^:DsRed) (n=9) embryos, showing a significantly increased phagocytosis level by microglial cells in the presence of hTau^P301L^-expressing neurons. (**E-H**, Video 4) Time-lapse image sequences from the optic tectum of a double transgenic Tg(ApoE-eGFP; HuC-hTau^P301L^:DsRed) 7 dpf embryo, showing a detail of a microglial cell in the process of phagocyting a neuron labelled with an apoptosis marker, acridine orange (merge: **E**, GFP and acridine: **F**, DsRed only: **G**, acridine only: **H**). The microglial cell filled with other dead tauopathic neurons extends its process to another dying tauopathic neuron and draws it towards its body cell to complete the phagocytosis process. (**I**, *p* = 0.027), Quantification of the number of non-engulfed apoptotic neurons in Tg(ApoE-eGFP; HuC-RFP) (n=11) and double transgenic Tg(ApoE-eGFP; HuC-hTau^P301L^:DsRed) (n=4) embryos in which there is a significantly higher number of non-engulfed apoptotic neurons. ***, *p* < 0.001; **, *p* < 0.01; *, *p* < 0.05. Scale bar (B, C, E, F, G, H) = 20 µm.

## 3 Discussion

To date, few studies have been conducted in *in vivo* conditions in healthy mice brains to show detailed morphological characterization of microglia (Cătălin et al., 2017; Sun et al., 2019). However, all studies aimed at investigating the physiology of microglia or their interactions with neurons in rodent models of neuronal diseases have relied widely on *ex vivo* and *in vitro* approaches, which cannot accurately reproduce the complexity of the physiological conditions observed in living brains (Bemiller et al., 2017; Hickman et al., 2013; Maphis et al., 2015; Rustenhoven et al., 2018).

While these marker-based approaches remain useful to gather prerequisite knowledge on immune cells, it is nonetheless crucial to preserve the morphology and dynamics of these highly plastic cells, which respond to very small changes in the CNS, and so to study them in a living brain (He et al., 2018). Recent studies show that time spent by microglia *ex vivo* is associated with a different evolution of gene expression until their expression levels become the reverse of the initial measures (Gosselin et al., 2017).

The present work is, to our knowledge, the first aimed at characterizing the dynamic behavior of microglial cells in the presence of pathological neurons expressing a human mutant Tau protein, hTau^P301L^, causing tauopathy.

Our results show that the presence of these hTau^P301L^-expressing neurons caused dramatic changes to microglia, with the cells displaying an amoeboid-like shape and higher mobility. Although these morphological and dynamic changes are reminiscent of the classical microglial activation profile seen in response to injury or disease (Nakajima and Kohsaka, 2001), these rounded microglial cells did not overexpress known pro-inflammatory cytokines, IL-1β and IL-8 showing that the observed changes were noninflammatory (Zhao et al., 2018). However, genetic depletion of microglia in brains containing hTau^P301L^-expressing neurons induced a markedly increased expression of both pro-inflammatory cytokines. This increased cytokine expression is reminiscent to that observed in a model of prion-induced neurodegeneration in mice (Zhu et al., 2016). One possible hypothesis is that astrocytes, the largest glial group, can also produce pro-inflammatory factors and exhibit a reactive state as it has been reported in tauopathy mice models (Sidoryk-Wegrzynowicz et al., 2017). This neuroinflammation could be exacerbated by the higher levels of pathological hyperphosphorylated Tau protein (Martini-Stoica et al., 2018; Perea et al., 2019).

In Tg(HuC-hTau^P301L^:DsRed) embryos, highly dynamic microglial cells displayed an intense phagocytic activity, specifically eliminating nearly twice as many apoptotic neurons as microglial cells in healthy brains. However, the significantly higher number of non-engulfed apoptotic neurons in tauopathic brains suggests that these microglial cells are overwhelmed by the excessive neuron death rate generated in this transgenic model. One therapeutic approach might thus be to enhance the phagocytic activity of microglia to slow the spread of the disease.

This study using intact zebrafish brain visualizes interactions between microglia and hTau^P301L^-expressing neurons in real time and sheds light on microglia activities exerting a protective role mainly through specific phagocytosis of apoptotic hTau^P301L^-expressing neurons, thereby limiting the spread of noxious cell bodies or pathologic hyperphosphorylated Tau. However, while displaying enhanced phagocytic activity towards hTau^P301L^-expressing neurons and efficiently eliminating dead neurons, microglial cells appeared overwhelmed, as evidenced by the higher number of dead, albeit non-engulfed dead neurons in transgenic embryo brains. These findings support therapeutic approaches based on the modulation of microglial phagocytic activity in a specific neurodegenerative context.

## 4 Materials and methods

### 4.1 Ethics statement

All the animal experiments described in the present study were conducted at the French National Institute of Health and Medical Research (INSERM) UMR 1141 in Paris in accordance with European Union guidelines for the handling of laboratory animals (http://ec.europa.eu/environment/chemicals/lab_animals/home_en.htm) and were approved by the Direction Départementale de la Protection des Populations de Paris and the French Animal Ethics Committee under reference No. 2012-15/676-0069.

### 4.2 Zebrafish lines and maintenance

Zebrafish were maintained at 26.5 °C in 14 h light and 10 h dark cycles. Embryos were collected by natural spawning and to avoid pigmentation, 0.003% 1-phenyl-2-thiourea (PTU) was added at 1 dpf (day post-fertilization). Transgenic Tg(HuC-hTau^P301L^:DsRed) embryos (Paquet et al., 2009), showing mosaic neuronal expression of hTau^P301L^ mutan protein, linked to FTDP-17, was used to reproduce key pathological features of tauopathy. In order to simultaneously observe microglia, we used the Tg(ApoE-eGFP) transgenic line (Peri and Nüsslein-Volhard, 2008) that allows live imaging of microglial cells with GFP. To investigate the consequences of the absence of microglia, we used the *nlrc3-like^st73/st73^* mutants (Shiau et al., 2013), in which the st73 recessive loss of function mutation in the noncanonical NOD-like receptor (NLR) gene is responsible for the absence of microglia in the brain.

### 4.3 Confocal imaging

For *in vivo* imaging, 7 dpf larvae were anaesthetized with 112 µg/ml 3-aminobenzoic acid ethyl ester (tricaine, Sigma), immobilized in 1.2% low melting-point agarose in the centre of a 35 mm glass-bottomed dish (Corning®), and covered with E3 medium containing 112 µg/ml tricaine. Images were acquired using a Leica SP8 confocal scanning laser microscope equipped with a Leica 20x/0.75 multi-immersion objective equipped with an Olympus 40x/1.1 water objective; or a Leica DM6000FS Spinning disk L2 microscope equipped with a Leica 25x/0.95 water immersion objective. All the images were then processed using LAS-X (Leica), MetaMorph 7.8.9 (Molecular Devices), AutoQuant X3.1.1 (Media Cybernetics), Fiji (Version 2.0.0-rc-65/1.52b) and Adobe Photoshop 7.0 (Adobe System).

### 4.4 Image analysis

The surface area, volume and sphericity 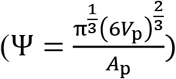 of microglial cells were quantified using Imaris MeasurementPro (Bitplane Inc.). The speed (distance travelled per unit time) and displacement (distance between first and last positions) of microglial processes were analysed using Imaris Filament tracer (Bitplane Inc.) on 15 minute long time-lapses. Microglial cell body displacements (distance between first and last positions) were tracked with Imaris MeasurementPro on 30 minute long time-lapses. Three-dimensional cell reconstructions were created using Imaris MeasurementPro.

### 4.5 Ablation of microglia

Morpholino pU.1 (MO-pU1): 5’-GATATACTGATACTCCATTGGTGGT-3’ designed to inhibit *pU1* mRNA translation, was obtained from Gene Tools. 2 nl of a 0.5 mM solution, corresponding to 1 pmol of pU.1 morpholino was injected into 1 to 2 cells stage embryos using standard protocols. After injection, the embryos were incubated in E3 medium at 28.5 °C until analysis at the desired stage. To select embryos in which microglia differentiation was fully blocked, Neutral Red staining was used to label microglia. Embryos were incubated in Neutral Red diluted in E3 medium for 5-8 hours at 28.5 °C, and rinsed 10 min before examination using a stereomicroscope (Zeiss).

### 4.6 Apoptosis labelling

To visualize apoptotic neurons, embryos were incubated in an Acridine Orange solution (1:500, VectaCell) for 20 min at 28.5 °C in the dark, and rinsed twice for 10 min in E3 medium. Although both GFP and acridine orange have very close excitation and emission spectra, their signals are easily distinguishable, with acridine orange emitting a much more intense fluorescence. Therefore, GFP channel (green) also shows Acridine Orange staining (blue) (Figure 3F).

**Figure 4.**
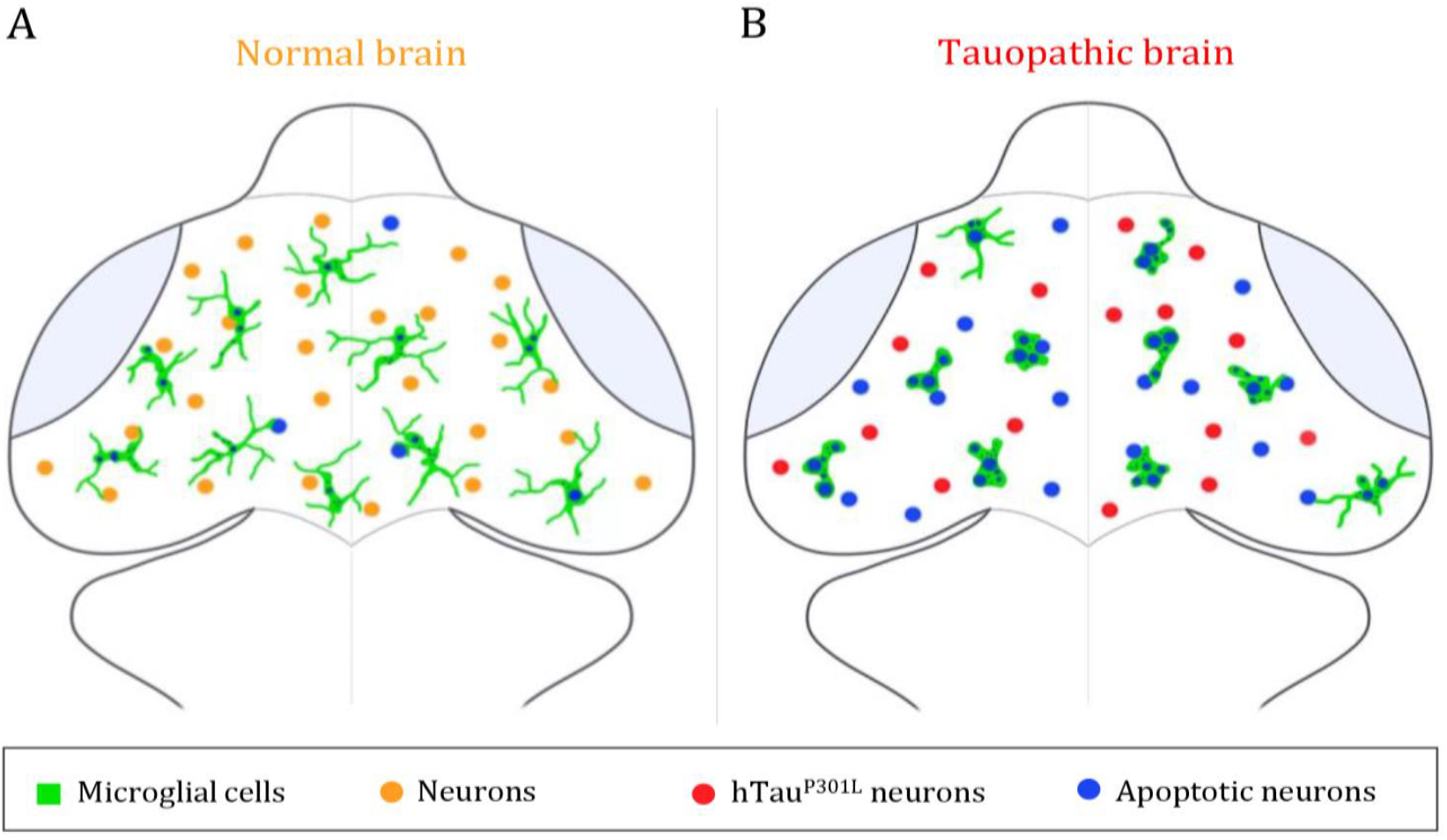
Summary illustration. (**A, B**) Brain illustrations of control embryo (**A**) and tauopathic embryo (**B**). In the control embryo brain, microglial cells (green) display a highly ramified morphology, allowing them to scan the brain and monitor neighbouring neurons (orange) and eliminate apoptotic ones (blue). However, in presence of hTau^P301L^–expressing neurons (red), microglial cells (green) adopt an amoeboid morphology, that allows them to move faster throughout the brain in order to eliminate tauopathic neurons undergoing apoptosis (blue). In spite of an increased phagocytic rate of microglial cells in the tauopathic brain, there is a higher number of non-engulfed apoptotic neurons (blue), in comparison to the control brain; thus, suggesting a saturated phagocytic capacity of microglial in the tauopathic brain.

### 4.7 Immunocytochemistry

6 dpf Tg(HuC-hTau^P301L^:DsRed; *nlrc3-like^st73/+^*) and Tg(HuC-hTau^P301L^:DsRed; *nlrc3-like^st73/73^*) embryos were anaesthetized in 0.2% tricaine, fixed with 4% paraformaldehyde, cryoprotected in 10% sucrose solution prior to flash freezing in isopentane. Samples were stored at −80°C until use. Embryos were cut into 20-µm-thick sections on cryostat, mounted on superfrost slides, and stored at −80°C. Cryosections (20 µm) were fixed in 4% paraformaldehyde at room temperature for 10 min. After washing thrice with PBS, sections were treated with 0.25% trypsin in 1X PBS for 2 min at 25°C. Immunohistochemistry was performed as previously described (Puverel et al., 2009). Briefly, sections were blocked and permeabilized with 0.2% gelatin, 0.25% Triton X-100 diluted in 1X PBS for 1 hr at room temperature and then incubated overnight at room temperature with anti-PHF1 (1:100, mouse monoclonal, gift of Dr. Peter Davies, Albert Einstein College of Medicine, New York, USA) After several washes, sections were incubated for 1 hr with the donkey anti-mouse coupled to Alexa Fluor 488 (1:500, Jackson Laboratories, West Grove, PA). Sections were counterstained for 10 min with 0.1% DAPI (Sigma-Aldrich) before being mounted with Vectashield Mounting Medium (Vector). Sections were analyzed using a Leica TCS SP8 confocal scanning system (Leica Microsystems). Images were acquired using a Leica SP8 confocal scanning laser microscope equipped with a Leica 20x/0.75 multi-immersion objective. Images were processed with LAS-X (Leica), Fiji (Version 2.0.0-rc-65/1.52b) and Adobe Photoshop 7.0 (Adobe System).

For whole mount immunostaining, 5 dpf wild-type embryos with or without microglia, were fixed in 4% formaldehyde in PBS for 1 hour 30 minutes at room temperature, washed three times in PBS (10 minutes each) and permeabilized in cold acetone (−20°C) for 20 minutes. After several washes, embryos were incubated in collagenase solution for 1 hour. Immunocytochemistry was performed as described previously (Naini et al., 2018) using rabbit anti-zebrafish L-plastin polyclonal antibody (gift of Dr. Michael Redd, University College London, United Kingdom), followed by Alexa-coupled secondary anti-rabbit antibody (Molecular Probes) at 1:500 dilution. After washing, the fluorescence was analyzed using a Leica TCS SP8 confocal scanning system (Leica Microsystems). Images were collected using a Leica 20x/0.75 multi-immersion objective. Images were processed with LAS-X (Leica), Fiji (Version 2.0.0-rc-65/1.52b) and Adobe Photoshop 7.0 (Adobe System).

### 4.8 RT-qPCR

Total RNAs were extracted from independent batches of 15 embryos each, using the NucleoSpin RNA kit (Macherey Nagel, Germany). Concentration of RNAs were assessed by spectrophotometry using a Nanodrop^TM^ device (Thermoscientific, USA). Total RNA (1µg) samples were reverse transcribed using the iScript^TM^ cDNA synthesis kit (Bio-Rad, USA). RT-qPCR experiments were performed in triplicate using SYBR Green Super-mix (Bio-Rad, USA) according to a program of 40 cycles in 3 steps (denaturation of 5 seconds at 96 ° C, hybridization of 10 seconds at 60 ° C and extension of 10 seconds at 72 ° C). Primers were designed manually following visual inspection of gene sequences. Gene sequences and NCBI references are given in **Supplementary table 1**. Specific mRNA levels were evaluated after normalization of the results with tubulin-α (tuba1) mRNA as reference, and the results were indicated in arbitrary units determined respectively to the levels of RNA determined in wild-type embryos and assessed using a Welch two-sample t-test or an ANOVA followed by a Tukey post-test.

### 4.9 Western blot

5 dpf embryos were collected, anaesthetized and lysed on ice with lysis buffer (50 mM Tris-HCl, 320mM Sucrose, pH 7.4) supplemented with protease and phosphatase inhibitors (Roche). Lysates were homogenized by sonication (thrice 10 seconds) and centrifuged at 600 g for 10 min. Samples containing 10 µg proteins were subjected to SDS-PAGE in 4-20% gradient acrylamide gel. Primary antibody against phosphorylated tau, Ser396 (1:1000, mouse monoclonal, Ozyme); and anti-human total tau antibody (1:1000, rabbit polyclonal, Dako Cytomation) were used. Subsequently, the blots were incubated for 1 hour at room temperature with the corresponding secondary antibodies (anti-mouse or anti-rabbit, 1:5000, Cell Signalling Technology) diluted in bovine serum albumin solution and developed with ECL RevelBlOt^®^ Plus (Ozyme) following manufacturer’s instructions. All statistics were assessed using a Welch two-sample t-test and all data are indicated as means ± SEM.

### 4.10 Statistics

All statistics were assessed using a Welch two-sample *t*-test or an ANOVA followed by a Tukey post-test. All data are represented as means ± SEM.

## Supporting information

Hassan-Abdi. R-Video 1

Hassan-Abdi. R-Video 2

Hassan-Abdi. R-Video 3

Hassan-Abdi. R-Video 4

Hassan-Abdi. R-Video 5

Hassan-Abdi. R-Video 6

Hassan-Abdi. R-Video 7

## 7 Acknowledgments

We thank Bettina Schmid (DZNE, Munich, Germany) for providing us with the Tg(HuC-hTau^P301L^:DsRed) transgenic line, Francesca Peri (University of Zurich, Zurich, Switzerland) for providing us with the Tg(ApoE-eGFP) and Michael Redd (University College London, London, United Kingdom) for providing us with the rabbit anti-zebrafish L-plastin polyclonal antibody. We also thank the imaging facility IMAG’IC (Cochin Institute, Paris). We also thank Christiane Romain and Olivier Bar (INSERM UMR 1141) for their technical assistance.

## Bibliography

A zebrafish model of tauopathy allows in vivo imaging of neuronal cell death and drug evaluation (2009). J. Clin. Invest. 119, 1382–1395. doi:10.1172/JCI37537.

Alavi Naini, S. M., and Soussi-Yanicostas, N. (2015). Tau Hyperphosphorylation and Oxidative Stress, a Critical Vicious Circle in Neurodegenerative Tauopathies? Oxid. Med. Cell. Longev. 2015. doi:10.1155/2015/151979.

Bemiller, S. M., McCray, T. J., Allan, K., Formica, S. V., Xu, G., Wilson, G., et al. (2017). TREM2 deficiency exacerbates tau pathology through dysregulated kinase signaling in a mouse model of tauopathy. Mol. Neurodegener. 12. doi:10.1186/s13024-017-0216-6.

Bhaskar, K., Konerth, M., Kokiko-Cochran, O. N., Cardona, A., Ransohoff, R. M., and Lamb, B. T. (2010). Regulation of Tau Pathology by the Microglial Fractalkine Receptor. Neuron 68, 19. doi:10.1016/j.neuron.2010.08.023.

Brockhaus, J., Möller, T., and Kettenmann, H. (1996). Phagocytozing ameboid microglial cells studied in a mouse corpus callosum slice preparation. Glia 16, 81–90. doi:10.1002/(SICI)1098-1136(199601)16:1<81::AID-GLIA9>3.0.CO;2-E.

Butovsky, O., and Weiner, H. L. (2018). Microglial signatures and their role in health and disease. Nat. Rev. Neurosci. doi:10.1038/s41583-018-0057-5.

Cătălin, B., Stopper, L., Bălşeanu, T.-A., and Scheller, A. (2017). The in situ morphology of microglia is highly sensitive to the mode of tissue fixation. J. Chem. Neuroanat. 86, 59–66. doi:10.1016/j.jchemneu.2017.08.007.

Eyo, U. B., and Dailey, M. E. (2013). Microglia: Key elements in neural development, plasticity, and pathology. J. Neuroimmune Pharmacol. 8, 494–509. doi:10.1007/s11481-013-9434-z.

Gosselin, D., Gosselin, D., Skola, D., Coufal, N. G., Holtman, I. R., Johannes, C. M., et al. (2017). An environment-dependent transcriptional network specifies human microglia identity. 3222, 33–35.

Hanisch, U. K. (2002). Microglia as a source and target of cytokines. Glia 40, 140–155. doi:10.1002/glia.10161.

Hanisch, U. K., and Kettenmann, H. (2007). Microglia: Active sensor and versatile effector cells in the normal and pathologic brain. Nat. Neurosci. 10, 1387–1394. doi:10.1038/nn1997.

Hansen, D. V., Hanson, J. E., and Sheng, M. (2018). Microglia in Alzheimer’s disease. J. Cell Biol. 217, 459–472. doi:10.1083/jcb.201709069.

He, Y., Yao, X., Taylor, N., Bai, Y., Lovenberg, T., and Bhattacharya, A. (2018). RNA sequencing analysis reveals quiescent microglia isolation methods from postnatal mouse brains and limitations of BV2 cells. J. Neuroinflammation 15. doi:10.1186/s12974-018-1195-4.

Hickman, S. E., Kingery, N. D., Ohsumi, T. K., Borowsky, M. L., Wang, L., Means, T. K., et al. (2013). The microglial sensome revealed by direct RNA sequencing. Nat. Neurosci. 16, 1896–1905. doi:10.1038/nn.3554.

Hu, X., Leak, R. K., Shi, Y., Suenaga, J., Gao, Y., Zheng, P., et al. (2015). Microglial and macrophage polarization - New prospects for brain repair. Nat. Rev. Neurol. 11, 56–64. doi:10.1038/nrneurol.2014.207.

Hutton, M., Lendon, C. L., Rizzu, P., Baker, M., Froelich, S., Houlden, H. H., et al. (1998). Association of missense and 5’-splice-site mutations in tau with the inherited dementia FTDP-17. Nature 393, 702–704. doi:10.1038/31508.

Kierdorf, K., and Prinz, M. (2017). Microglia in steady state. J. Clin. Invest. 127, 3201–3209. doi:10.1172/JCI90602.

Laurent, C., Buée, L., and Blum, D. (2018). Tau and neuroinflammation: What impact for Alzheimer’s Disease and Tauopathies? Biomed. J. 41, 21–33. doi:10.1016/j.bj.2018.01.003.

Leong, S. -K, and Ling, E. -A (1992). Amoeboid and ramified microglia: Their interrelationship and response to brain injury. Glia 6, 39–47. doi:10.1002/glia.440060106.

Ling, E. -A, and Wong, W. -C (1993). The origin and nature of ramified and amoeboid microglia: A historical review and current concepts. Glia 7, 9–18. doi:10.1002/glia.440070105.

Maphis, N., Xu, G., Kokiko-cochran, O. N., Cardona, A., Ransohoff, R. M., Lamb, B. T., et al. (2015a). Loss of tau rescues inflammation-mediated neurodegeneration. Front. Neurosci. 9, 1–12. doi:10.3389/fnins.2015.00196.

Maphis, N., Xu, G., Kokiko-Cochran, O. N., Jiang, S., Cardona, A., Ransohoff, R. M., et al. (2015b). Reactive microglia drive tau pathology and contribute to the spreading of pathological tau in the brain. Brain 138, 1738–1755. doi:10.1093/brain/awv081.

Martini-Stoica, H., Cole, A. L., Swartzlander, D. B., Chen, F., Wan, Y.-W., Bajaj, L., et al. (2018). TFEB enhances astroglial uptake of extracellular tau species and reduces tau spreading. J. Exp. Med. 215, 2355–2377. doi:10.1084/JEM.20172158.

Naini, S. M. A., Yanicostas, C., Hassan-Abdi, R., Blondeel, S., Bennis, M., Weiss, R. J., et al. (2018). Surfen and oxalyl surfen decrease tau hyperphosphorylation and mitigate neuron deficits in vivo in a zebrafish model of tauopathy. Transl. Neurodegener. 7, 6. doi:10.1186/s40035-018-0111-2.

Nakajima, K., and Kohsaka, S. (2001a). Microglia: Activation and their significance in the central nervous system. J. Biochem. doi:10.1093/oxfordjournals.jbchem.a002969.

Nakajima, K., and Kohsaka, S. (2001b). Microglia: Activation and their significance in the central nervous system. Oxford University Press doi:10.1093/oxfordjournals.jbchem.a002969.

Nimmerjahn, A., Kirchhoff, F., and Helmchen, F. (2005). Resting microglial cells are highly dynamic surveillants of brain parenchyma in vivo. Science (80-.). 308, 1314–1318. doi:10.1126/science.1110647.

Paquet, D., Bhat, R., Sydow, A., Mandelkow, E. M., Berg, S., Hellberg, S., et al. (2009). A zebrafish model of tauopathy allows in vivo imaging of neuronal cell death and drug evaluation. J. Clin. Invest. 119, 1382–1395. doi:10.1172/JCI37537.

Perea, J. R., Llorens-Martín, M., Ávila, J., and Bolós, M. (2018). The Role of Microglia in the Spread of Tau: Relevance for Tauopathies. Front. Cell. Neurosci. 12. doi:10.3389/fncel.2018.00172.

Perea, J. R., López, E., Díez-Ballesteros, J. C., Ávila, J., Hernández, F., and Bolós, M. Extracellular Monomeric Tau Is Internalized by Astrocytes. Front. Neurosci. doi:10.3389/fnins.2019.00442.

Peri, F., and Nüsslein-Volhard, C. (2008). Live Imaging of Neuronal Degradation by Microglia Reveals a Role for v0-ATPase a1 in Phagosomal Fusion In Vivo. Cell 133, 916–927. doi:10.1016/j.cell.2008.04.037.

Puverel, S., Nakatani, H., Parras, C., and Soussi-Yanicostas, N. (2009). Prokineticin receptor 2 expression identifies migrating neuroblasts and their subventricular zone transient-amplifying progenitors in adult mice. J. Comp. Neurol. 512, 232–242. doi:10.1002/cne.21888.

Rhodes, J., Hagen, A., Hsu, K., Deng, M., Liu, T. X., Look, A. T., et al. (2005). Interplay of pu.1 and Gata1 determines myelo-erythroid progenitor cell fate in zebrafish. Dev. Cell 8, 97–108. doi:10.1016/j.devcel.2004.11.014.

Rustenhoven, J., Smith, A. M., Smyth, L. C., Jansson, D., Scotter, E. L., Swanson, M. E. V., et al. (2018). PU.1 regulates Alzheimer’s disease-associated genes in primary human microglia. Mol. Neurodegener. 13. doi:10.1186/s13024-018-0277-1.

Shiau, C. E., Monk, K. R., Joo, W., and Talbot, W. S. (2013). An Anti-inflammatory NOD-like Receptor Is Required for Microglia Development. Cell Rep. 5, 1342–1352. doi:10.1016/j.celrep.2013.11.004.

Sidoryk-Wegrzynowicz, M., Gerber, Y. N., Ries, M., Sastre, M., Tolkovsky, A. M., and Spillantini, M. G. (2017). Astrocytes in mouse models of tauopathies acquire early deficits and lose neurosupportive functions. Acta Neuropathol. Commun. 5, 89. doi:10.1186/s40478-017-0478-9.

Sousa, C., Golebiewska, A., Poovathingal, S. K., Kaoma, T., Pires-Afonso, Y., Martina, S., et al. (2018). Single-cell transcriptomics reveals distinct inflammation-induced microglia signatures. EMBO Rep. doi:10.15252/embr.201846171.

Spillantini, M. G., and Goedert, M. (2013). Tau pathology and neurodegeneration. Lancet Neurol. 12, 609–622. doi:10.1016/S1474-4422(13)70090-5.

Sun, W., Suzuki, K., Toptunov, D., Stoyanov, S., Yuzaki, M., Khiroug, L., et al. (2019). In vivo Two-Photon Imaging of Anesthesia-Specific Alterations in Microglial Surveillance and Photodamage-Directed Motility in Mouse Cortex. Front. Neurosci. 13, 421. doi:10.3389/fnins.2019.00421.

Thameem Dheen, S., Kaur, C., and Ling, E.-A. (2007). Microglial Activation and its Implications in the Brain Diseases. Curr. Med. Chem. 14, 1189–1197. doi: http://dx.doi.org/10.2174/092986707780597961.

Wake, H., Moorhouse, A. J., Miyamoto, A., and Nabekura, J. (2013). Microglia: Actively surveying and shaping neuronal circuit structure and function. Trends Neurosci. 36, 209–217. doi:10.1016/j.tins.2012.11.007.

Wang, Y., and Mandelkow, E. (2016). Tau in physiology and pathology. Nat. Rev. Neurosci. 17, 5–21. doi:10.1038/nrn.2015.1.

Zhao, X., Liao, Y., Morgan, S., Mathur, R., Feustel, P., Mazurkiewicz, J., et al. (2018). Noninflammatory Changes of Microglia Are Sufficient to Cause Epilepsy. Cell Rep. 22, 2080–2093. doi:10.1016/j.celrep.2018.02.004.

Zhu, C., Herrmann, U. S., Falsig, J., Abakumova, I., Nuvolone, M., Schwarz, P., et al. (2016). A neuroprotective role for microglia in prion diseases. J. Exp. Med. 213, 1047–59. doi:10.1084/jem.20151000.

