## Supplementary figures and images for "Neurons expressing pathological Tau protein trigger dramatic changes in microglial morphology and dynamics"

### Supplementary Figure 1

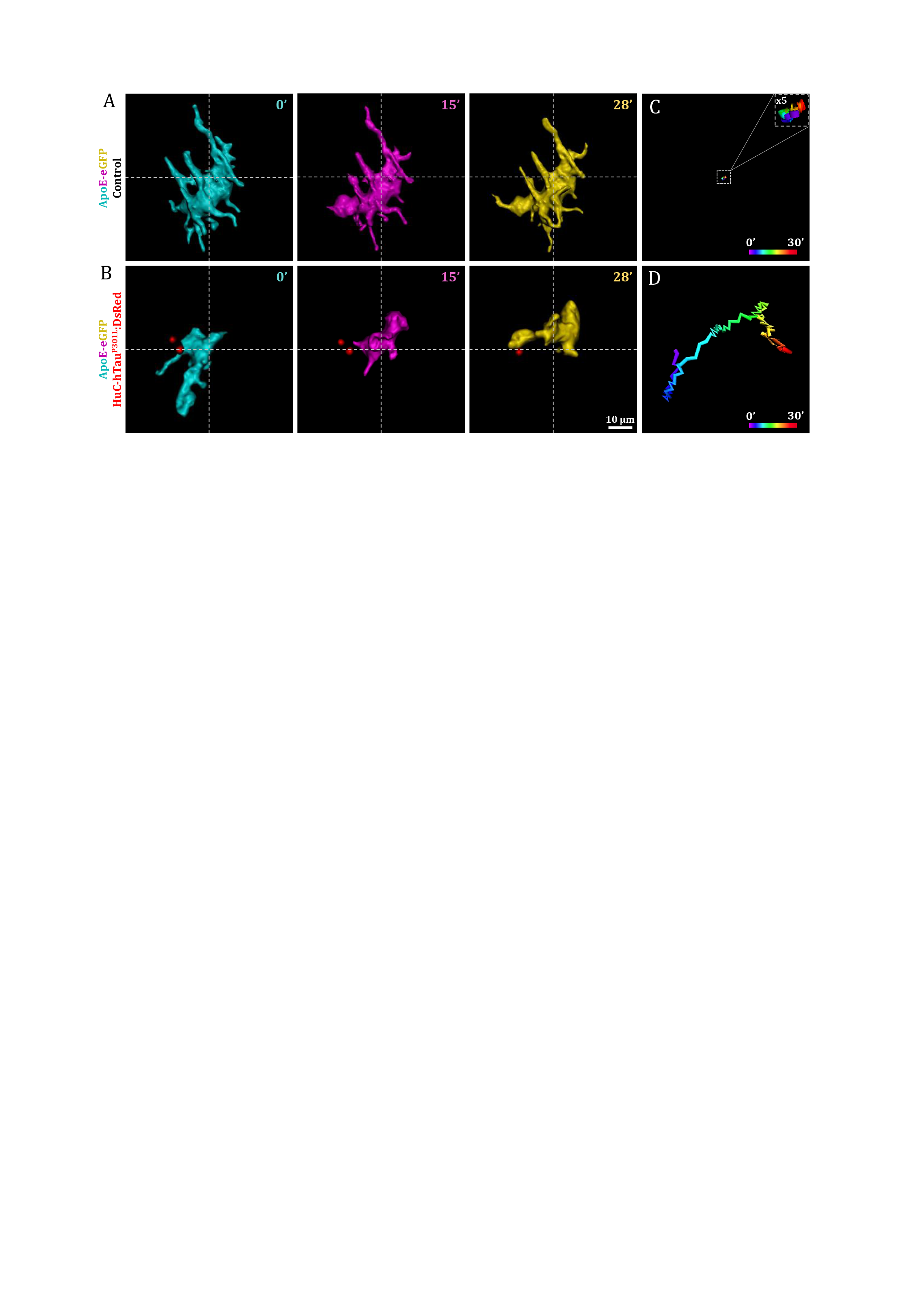

### supplementary Figure 2

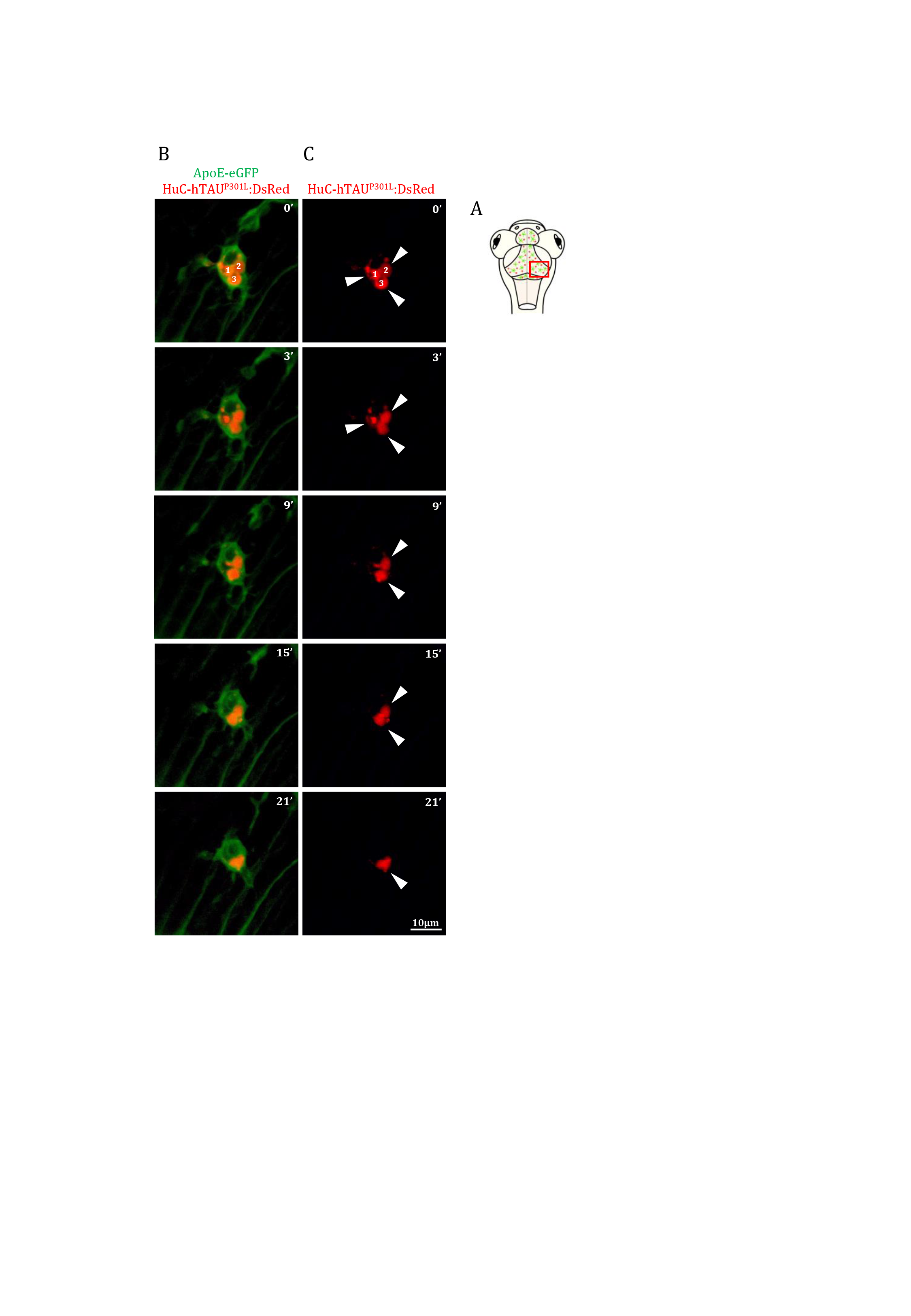
