## supplementary Material for "Neurons expressing pathological Tau protein trigger dramatic changes in microglial morphology and dynamics"

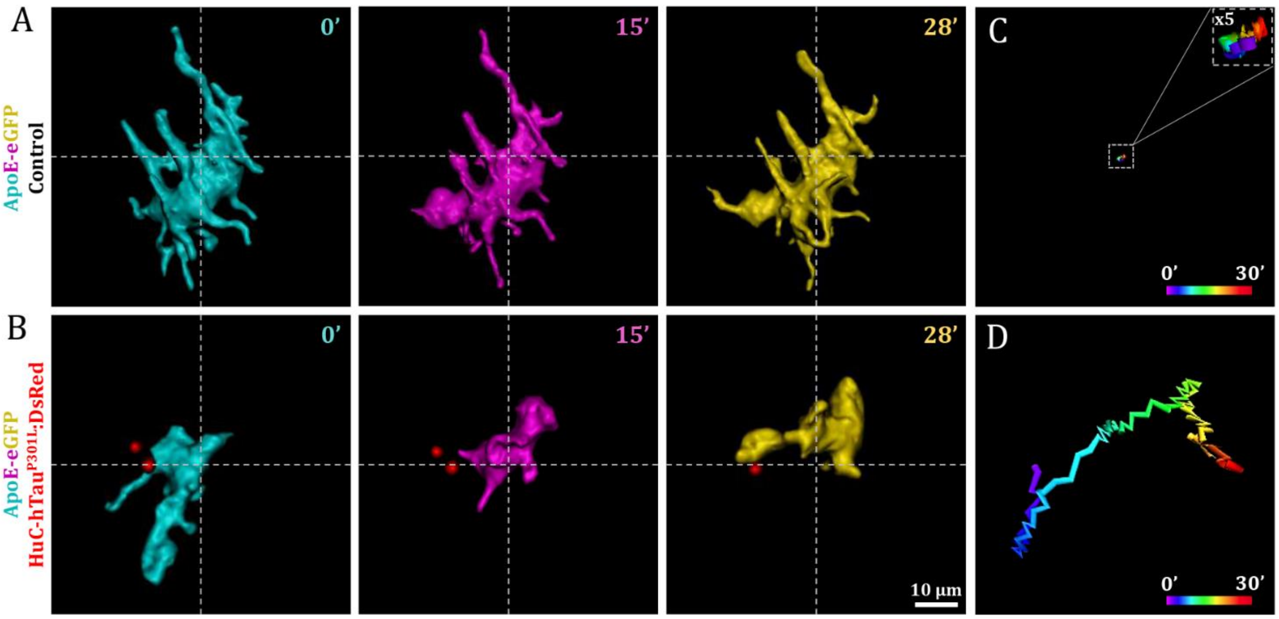


**Supplementary figure 1. 3D reconstruction of time-lapse showing microglia mobility.** (**A-D**) Three-dimensional reconstruction of time-lapse sequences showing the details of the mobility of microglial cell bodies (**A, B**) and the corresponding tracking of cell body displacement (**C, D**) in Tg(ApoE-eGFP) (**A, C**) and Tg(ApoE-eGFP; HuC-hTau$^{P301L}$:DsRed) embryos (**B, D**). The microglial cell from the wild-type brain (**A**) displays very restricted cell body movement as shown by the tracking (**C**) compared to the highly mobile microglial cells (**B**) found next to hTau$^{P301L}$-expressing neurons (red) (**D**). Related to videos 5 and 6. Scale bar (**A, B**) = 10 µm.

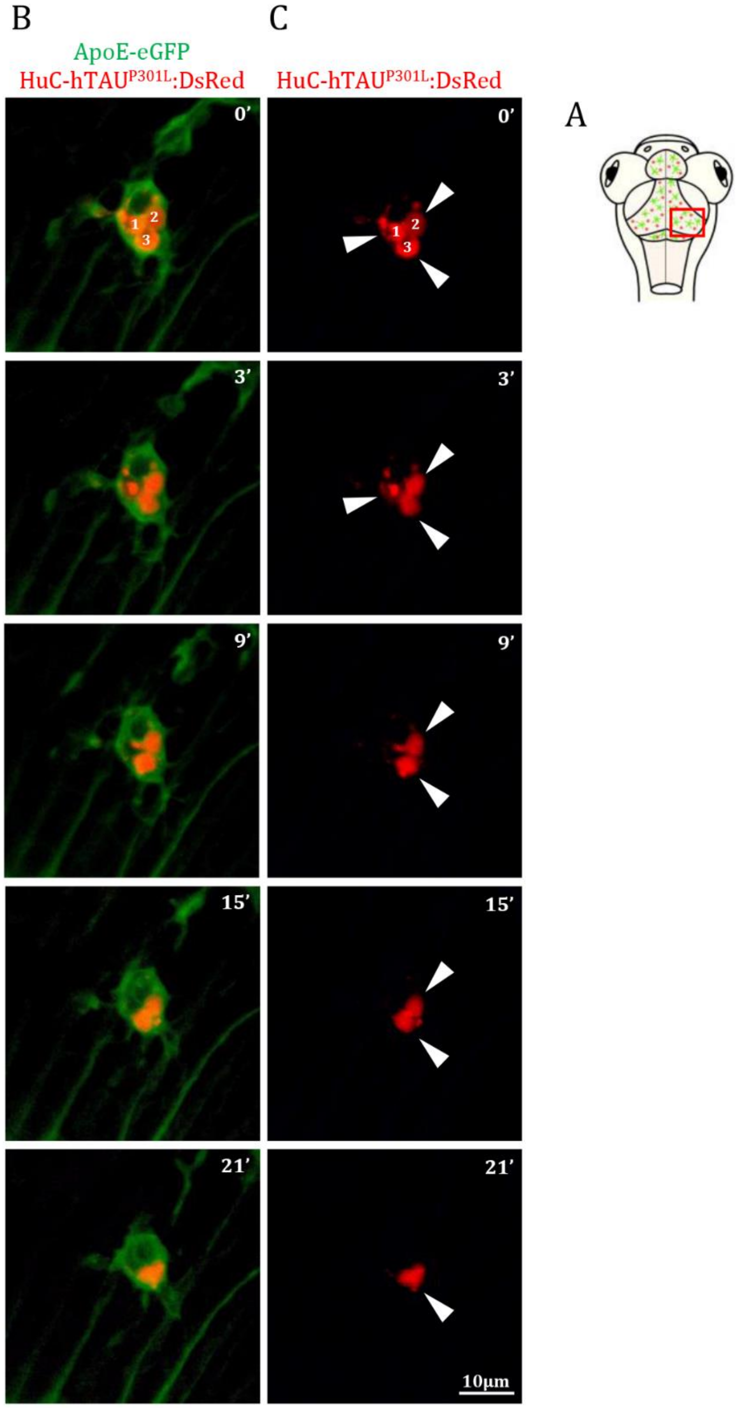


**Supplementary figure 2. Phagocytosis of diseased neurons by microglia *in vivo*.** (**A**) Schematic illustration of 7 dpf embryo in dorsal view. The red square shows the region of the optic tectum where the time-lapse (**B, C**) was recorded. (**B, C**) Time-lapse image sequences from the optic tectum of a 7 dpf transgenic Tg(ApoE-eGFP; HuC-hTau$^{P301L}$:DsRed) embryo, showing a microglial cell phagocyting three diseased neurons and showing complete digestion of two of them after 20 minutes. Related to video 7. Scale bar (**B, C**) = 10 µm.

**Supplementary table 1. Sequence of primers used for RT-qPCR**

| Gene | Accession number | Nucleotide sequence (5' - 3’) |
|---|---|---|
| ***tuba-1*** | ENSDARG00000001889 | **Forward:** GAGCCCACTGCTATTGATGAG |
| | | **Reverse:** GTGCACTGGTCAGACAGTTTG |
| ***IL-1β*** | ENSDARG00000098700 | **Forward:** GCTGGAGATCCAAACGGATA |
| | | **Reverse:** ATACGCGGTGCTGATAAACC |
| ***IL-8*** | ENSDART00000161996.2 | **Forward:** TGACCATCATTGAAGGAATGAG |
| | | **Reverse:** CATCAAGGTGGCAATGATCTC |

**Supplementary videos**

**Video 1. Microglial dynamics in wild-type Tg(ApoE-eGFP) brain. Related to Figure 1H.** Time-lapse video of a dorsal view of the optic tectum of a 7 dpf Tg(ApoE-eGFP) embryo during 15 minutes, showing the dynamics of microglia in a wild-type brain, which mainly use their processes to monitor neighbouring cells. Time interval between frames: 30 s. Video speed: 8 frames/s.

**Video 2. Microglial dynamics in Tg(ApoE-eGFP; HuC-hTau$^{P301L}$:DsRed)**. **Related to Figure 1I.** Time-lapse video of a dorsal view of the optic tectum of a 7 dpf Tg(ApoE-eGFP; HuC-hTau$^{P301L}$:DsRed) embryo during 15 minutes, showing the increased dynamics of microglial cells in the presence of hTau$^{P301L}$-expressing neurons (DsRed labelling not shown), where microglial cells move over long distances. Time interval between frames: 30 s. Video speed: 8 frames/s.

**Video 3. Phagocytosis of a hTau$^{P301L}$-expressing neuron in Tg(ApoE-eGFP; HuC-hTau$^{P301L}$:DsRed) brain. Related to Figure 3B, C.** Time-lapse video of the dorsal view of the optic tectum of a 7 dpf Tg(ApoE-eGFP; HuC-hTau$^{P301L}$:DsRed) embryo during 30 minutes, showing the phagocytosis of a pathological hTau$^{P301L}$–expressing neuron (DsRed, white arrow) by a microglial cell (GFP) that extends a process to capture the neuron, and draw it towards its body cell to execute the complete engulfment of the neuron. Time interval between frames: 30 s. Video speed: 8 frames/s.

**Video 4. Phagocytosis of an apoptotic hTau$^{P301L}$–expressing neuron in Tg(ApoE-eGFP; HuC-hTau$^{P301L}$:DsRed) brain. Related to Figure 3E-H.** Time-lapse video of a microglial cell from the optic tectum of a 7 dpf Tg(ApoE-eGFP; hTau$^{P301L}$:DsRed) embryo, labelled with acridine orange, during 39 minutes, showing the phagocytosis of an apoptotic hTau$^{P301L}$–expressing neuron (white) by a microglial cell (green). Time interval between frames: 30 s. Video speed: 8 frames/s.

**Video 5. Detail of microglial mobility in wild-type Tg(ApoE-eGFP) brain. Related to Supplementary figures 1A, C.** Three-dimensional reconstruction of a time-lapse video of a microglial cell in the optic tectum of a 7 dpf Tg(ApoE-eGFP) embryo during 27 min, with a transparent 3D surface and tracking over time. The microglial cell body (green) is stationary as represented by the tracking line over time that is restrained at the centre of the cell body. Time interval between frames: 20 s. Video speed: 10 frames/s.

**Video 6. Detail of microglial mobility in Tg(HuC-hTau$^{P301L}$:DsRed) brain. Related to Supplementary figures 1B, D.** Three-dimensional reconstruction of a time-lapse video of a microglial cell in the optic tectum of a 7 dpf Tg(HuC-hTau$^{P301L}$:DsRed) embryo during 34 min, with a transparent 3D surface and tracking over time. The microglial cell body (green) migrates and changes position as represented by the tracking line that is spreading and interacting with hTau$^{P301L}$–expressing neurons (red). Time interval between frames: 30 s. Video speed: 10 frames/s.

**Video 7. Detail of phagocytosis of a hTau$^{P301L}$–expressing neuron in Tg(ApoE-eGFP; HuC-hTau$^{P301L}$:DsRed) brain. Related to Supplementary figure 2.** Time-lapse video of a microglial cell from the optic tectum of a 7 dpf Tg(ApoE-eGFP; HuC-hTau$^{P301L}$:DsRed) during 21 min, showing the phagocytosis of three hTau$^{P301L}$–expressing neurons (red, white arrow heads) by a microglial cell (green), with the complete digestion of two of them. Time interval between frames: 30 s. Video speed: 4 frames/s.
